# Differential modulation of temporal precision and perceived timing by tactile spatial and temporal context

**DOI:** 10.64898/2026.07.26.740772

**Authors:** Mehdi Adibi

## Abstract

Perception of sensory events depends not only on the stimulus itself but also on the surrounding spatial and temporal context. While contextual interactions have been extensively studied in vision, less is known about how tactile spatial and temporal context shape temporal representation of tactile landmarks embedded within continuous tactile signals. we investigated how neighbouring tactile stimuli influence the temporal precision and perceived timing of a target tactile landmark using an auditory-referenced temporal order judgement task. The number, location, and relative timing of task-irrelevant vibrotactile flankers were manipulated independently. Increasing the number of flankers reduced temporal precision, with comparable effects for adjacent and contralateral stimulation, indicating that temporal precision is limited by contextual interactions beyond local spatial proximity. In contrast, the relative timing of the flankers did not alter temporal sensitivity but systematically shifted perceived target timing. Flankers preceding the target shifted the perceived timing of the target earlier, whereas lagging flankers shifted it to the opposite direction. When one flanker preceded and the other followed the target the direction of the resulting spatiotemporal sequence across the fingers determined the direction of the perceptual bias. These findings reveal two distinct forms of contextual modulation in tactile temporal perception. Spatial context degrades temporal precision, whereas temporal context shifts perceived event timing. These findings demonstrate that spatial and temporal context shape distinct temporal aspects of tactile representations, extending contextual interference from feature identity and localisation to temporal precision and perceived timing.

## Introduction

Perception is shaped not only by the sensory signal of interest but also by the surrounding spatial and temporal context in which that signal occurs [1–3].Neighbouring stimuli in time and space influence the perceived properties of a target, including its apparent intensity [4], orientation [5, 6], size [7–9], position [10, 11], duration [12] and motion Addams1834, demonstrating that perception reflects interactions among sensory inputs rather than solely their independent processing [13]. These contextual interactions have been extensively studied in vision, where phenomena such as forward and backward masking [14, 15], crowding [16–19], uncrowding [20, 21], surround suppression [22, 23], contextual distortions [24, 25], and serial dependence [26] have been used to characterise how information from neighbouring locations or times is integrated, pooled, or suppressed. Comparable contextual effects have also been characterised through temporal masking [27], informational masking [28, 29], and the grouping and segregation of sounds into auditory streams and sources [30–33].

Touch likewise exhibits contextual interactions across both the skin surface and time. Tactile stimuli can be obscured by preceding, simultaneous, or subsequent stimulation, with interference depending on stimulus timing, frequency, and spatial arrangement [34–38]. Inputs delivered to different fingers can also suppress, compete with, or be integrated with one another [39–42]. The spatiotemporal arrangement of tactile signals delivered to different skin locations can transform their percept from independent events into a single localised sensation, an apparent tactile trajectory, or continuous motion across the skin and fingers [10, 43–46]. Together, these examples show that tactile context can both distort the representation of individual stimuli and construct coherent percepts extending across multiple stimulation sites, and that tactile features, such as perceived intensity, localisation, and motion, are dynamically computed relative to adjacent spatial and temporal context rather than in isolation.

These interactions are relevant during natural haptic exploration. Objects and surfaces typically extend beyond a single fingerpad. Relative motion between the fingers and object generates continuously evolving tactile signals [47]. Repeated exploratory movements provide successive samples of a surface, while simultaneous contact with several fingers distributes related information across multiple receptor populations. During exploration, information sampled by different fingerpads must be integrated into a coherent representation. Studies suggest multidigit tactile signals are integrated into an overall estimate of tactile motion, although their contribution depends on the spatial arrangement of the stimulated fingers and the direction of motion across the hand [41, 42]. In particular, multidigit motion signals are integrated rather than represented as separate estimates, while within-hand integration assigns greater perceptual weight to the finger leading the motion trajectory [41, 42].

Constructing a coherent tactile percept requires more than collecting information from a larger skin area. Sensory signals arriving at different fingers and at different moments must be related over time. This temporal organisation allows sequential or spatially distributed inputs to be interpreted as samples of the same surface or as components of a single motion trajectory [47]. It also supports perceptual stability despite changes in the temporal structure of peripheral input during exploration. For example, temporal patterns evoked in tactile afferents by fine textures expand or contract with scanning speed, whereas perceived texture remains stable [48, 49]. This speed invariance implies that the nervous system transforms temporally varying peripheral responses into a relatively stable representation of surface identity [49]. Relative timing between tactile signals also provides a direct cue for tactile motion: phase-shifted stimulation across fingers can generate a robust sense of motion even when the stimulated surfaces themselves remain stationary [44–46, 50].

Despite the central role of temporal relationships in tactile perception in shaping tactile percepts, little is known about the temporal precision with which an event embedded within a continuous tactile signal is represented when neighbouring fingers or skin locations are simultaneously stimulated [44, 48, 50–52]. Whether contextual tactile signals influence temporal precision, perceived timing, or both remains unknown. Previous studies have largely examined the temporal order of brief, discrete tactile stimuli delivered to different fingers, hands, or body locations [44, 53–57]. These studies have established that tactile temporal-order judgements depend on both somatotopic and external spatial representations and can be strongly disrupted by changes in limb posture [53, 54, 56, 58–62]. How spatial and temporal contextual signals independently influence temporal precision and perceived timing, however, remains unresolved.

Temporal order judgements provide a sensitive measure of perceived event timing and temporal sensitivity [55]. Temporal-order judgements provide a sensitive measure of perceived event timing and temporal precision [55]. Here, we used an auditory reference to estimate the perceived timing of a salient landmark embedded within a continuous amplitude-modulated tactile signal delivered to one finger. Unlike conventional tactile temporal-order tasks, which compare the onset of two discrete tactile events, the temporal-order judgement here measured the timing assigned to an internal landmark within an extended tactile signal. Task-irrelevant tactile flankers varied independently in number, spatial arrangement, and relative timing, enabling us to dissociate how spatial context influences temporal precision from how temporal context shifts perceived event timing.

The present study shows that increasing the number of flankers reduced temporal sensitivity irrespective of their spatial location, whereas the relative timing of the flankers systematically shifted perceived target timing without affecting temporal precision. These findings reveal dissociable influences of spatial and temporal context on the temporal representation of tactile events, selectively affecting temporal precision and perceived timing.

## Materials and methods

Psychophysical experiments were conducted to quantify how the spatial and temporal context of neighbouring tactile stimuli influences temporal perception. Participants performed a two-alternative forced-choice (2AFC) temporal order judgement (TOJ) task in which they judged whether the perceived peak of a target vibrotactile stimulus occurred before or after a brief auditory reference. The spatial configuration, number, and relative timing of task-irrelevant tactile flankers were systematically varied to dissociate their effects on temporal sensitivity and perceptual bias. Experimental conditions were presented in pseudorandom order, and participants completed self-paced trials throughout each session. All experimental procedures were approved by the Monash University Human Research Ethics Committee (MUHREC; Project ID 27649; approved 2 August 2021).

### Participants

A total of 10 healthy adults (8 female; age range 18–49 years; mean age 24 years, standard deviation (SD) 9.4 years; all right-handed) participated in the study. Participants were recruited from Monash University undergraduate and graduate students and the surrounding community. All participants reported normal tactile perception and no history of neurological disorders, and provided written informed consent prior to participation.

### Apparatus and Vibrotactile Stimulation

The apparatus and stimulation procedures were identical to those described previously [45, 46]. Briefly, vibrotactile stimuli were delivered simultaneously to fingertips using miniature solenoid transducers (PMT-20N12AL04-04, Tymphany HK Ltd; 4 Ω, 1 W, 20 mm diameter) mounted 5 cm apart on a vibration-isolated platform. Stimuli were generated in MATLAB (MathWorks Inc.) at a sampling rate of 44.1 kHz, and delivered a Creative Sound Blaster Live! 24-bit External (model SB0490). The shape and curvature of the transducer matched the size and contour of adult fingertips [63]. The solenoid transducers were driven using a 3 W stereo amplifier module (PAM8403 chipset). Solenoid transducer outputs were matched across channels and configurations. Sound pressure levels (SPL) were measured 5 mm from each actuator and adjusted to produce matched outputs (mean SPL = 78.9 dB). Matching was further verified perceptually to ensure comparable stimulus intensity across actuators.

Vibrotactile stimuli consisted of a sinusoidal carrier (*f_c_* = 100*Hz*), with the amplitude modulation, spatial configuration, and temporal parameters depended on the specific experimental task, as described below. Although the carrier frequency falls within the audible range, pilot testing confirmed that the stimuli were not audible to participants and were perceived only through touch [45]. Additionally, participants wore a headset blocking ambient auditory noise. The modulation amplitude was set well above detection threshold.

### Tactile Landmark Temporal Order Judgement Task

Participants performed a two-alternative forced-choice (2AFC) temporal order judgement (TOJ) task in which they judged whether a brief auditory probe occurred before or after the perceived peak of a target vibrotactile stimulus delivered to the middle fingertip of the right hand.

The target stimulus consisted of a single cycle of sinusoidal amplitude modulation (2 s duration; 0.5 Hz envelope frequency) with a 100 Hz carrier vibration. The target envelope was generated by raising the sinusoidal modulation to the fifth power (cos^5^), producing a single well-defined tactile landmark. Task-irrelevant flanker stimuli were presented simultaneously using broader cos^2^ envelopes. Depending on the experimental condition, no flanker was presented, or flankers were delivered to the right index finger, the right index and ring fingers, or the left index finger. For the flanker conditions, the envelope peak occurred either 60° before or 60° after the target peak. In the two-flanker condition, the two flankers were assigned independently, allowing both flankers to lead, both to lag, or one to lead while the other lagged the target.

The auditory reference was a 10 ms, 1 kHz tone presented binaurally through headphones. It occurred at signed temporal offsets of *±*50, *±*150, and *±*300 ms relative to the tactile landmark. Positive offsets corresponded to presentation of the auditory probe after the tactile landmark, whereas negative offsets corresponded to presentation before the tactile landmark. Participants indicated whether the auditory probe occurred before or after the perceived tactile peak using the left and right arrow keys.

Experimental conditions were presented in pseudorandom order. Each participant completed 25 repetitions of every combination of temporal offset and flanker configuration, resulting in 600 trials per participant and a total of 6,000 trials across the experiment.

### Psychometric Modelling

Trial-level responses were analysed using hierarchical generalised linear mixed-effects models (GLMMs) implemented in MATLAB (MathWorks Inc.). Two classes of models were fitted. Temporal discrimination accuracy was analysed using a binomial GLMM with a logit link, whereas target-first responses (psychometric choice functions) were analysed using a binomial GLMM with a probit link unless otherwise stated. Fixed effects represented the experimental variables under investigation (e.g., flanker number, spatial configuration, and relative flanker timing), while random effects accounted for between-participant variability. Subject-specific random intercepts and/or slopes were included where appropriate, and candidate random-effects structures were compared to identify the most parsimonious model.

For the accuracy models, discrimination performance was expressed as a function of the absolute temporal separation between the auditory reference and tactile landmark. Models were parameterised without an intercept, constraining performance to chance (50%) at zero temporal separation, consistent with the symmetry of the temporal order judgement task. Temporal sensitivity was quantified by the fitted slope parameter (*β*), with larger values corresponding to steeper psychometric functions and greater temporal precision.

Psychometric choice functions were modelled as the probability of reporting that the tactile landmark occurred before the auditory reference. Under the cumulative Gaussian assumption of the probit model, temporal uncertainty was estimated as *σ* = 1*/β*, where *β* denotes the fitted probit slope parameter. The point of subjective simultaneity (PSS) was estimated as PSS = *α/β*, where *α* and *β* denote the fitted intercept and slope, respectively. Standard errors (SE) for both *σ* and PSS were estimated using first-order error propagation (delta method) based on the covariance matrix of the fitted fixed-effect parameters.

Candidate models differing in their fixed- and random-effects structures were compared using the Akaike Information Criterion (AIC) and Bayesian Information Criterion (BIC). Where models were nested, likelihood-ratio tests were additionally used to quantify whether the increased model complexity provided a significantly better description of the data. Unless otherwise stated, the model with the lowest information criteria and the simplest parameterisation was used for subsequent analyses.

## Results

Perception requires extracting the timing of behaviourally relevant sensory input while ignoring competing stimuli. To investigate how adjacent tactile events influence temporal perception, participants performed a tactile temporal order judgement (TOJ) task in which they judged whether the peak of an amplitude-modulated vibration delivered to the middle finger occurred before or after a brief auditory reference of 10 ms (Figure 1A). Task-irrelevant vibrotactile flankers were presented either to one adjacent finger, to both adjacent fingers, or to the contralateral index finger, with flanker peaks occurring before or after the target (Figure 1B). This design allowed us to independently quantify how the number, spatial configuration, and temporal relationship of flankers influence temporal precision and perceived event timing.

**Figure 1.**
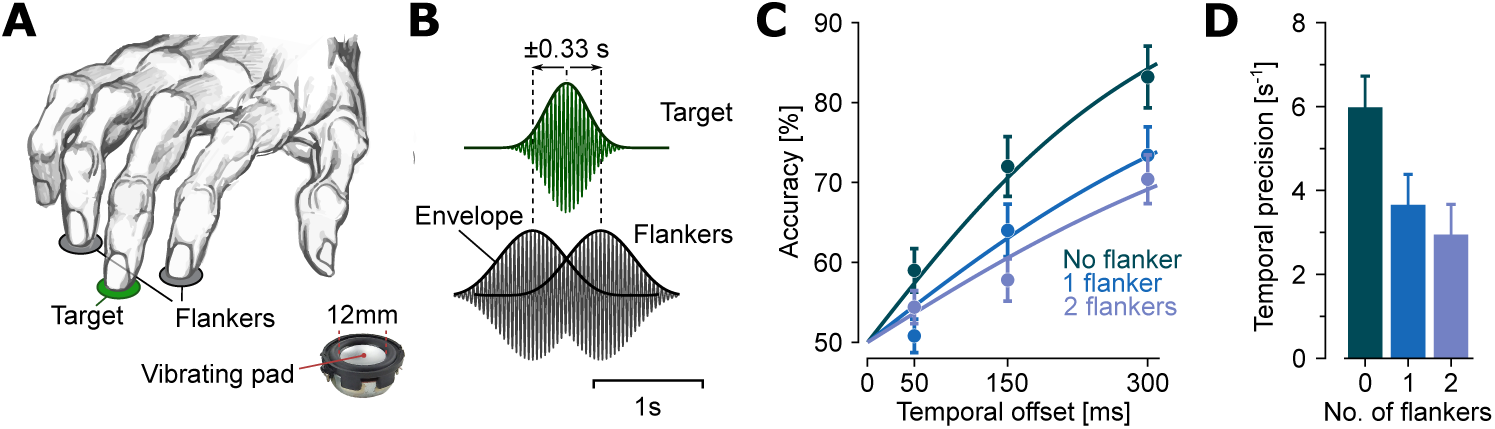
Experimental paradigm and the effect of number of flankers on accuracy and temporal precision. **A**, Schematic of the tactile temporal order judgement (TOJ) task. Participants judged whether the peak of the target vibration delivered to the middle finger occurred before or after a brief auditory reference while ignoring one or two flanking vibrotactile stimuli delivered to the adjacent fingers. The inset illustrates the miniature solenoid transducer used for vibrotactile stimulation. **B**, Target and flanker vibrotactile waveforms with their corresponding amplitude envelopes (bold curves). For illustration purposes, the carrier frequency is shown as 25 Hz to improve visibility, whereas all experiments were performed using a 100 Hz carrier. Flanker peaks either lead or lag the target peak by 0.33 s (60° phase difference at the modulation frequency of 0.5 Hz), with equal probability. **C**, Temporal order discrimination accuracy as a function of target–auditory reference temporal offset for the no-flanker, one-flanker, and two-flanker conditions. Markers represent the mean percentage of correct responses across participants. Error bars denote standard errors (SE) across participants. Solid curves show population-level predictions of the hierarchical generalised linear mixed-effects model. **D**, Model-derived estimates of temporal precision for each flanker condition obtained from the hierarchical generalised linear mixed-effects model. Error bars represent model-estimated standard errors.

### Temporal precision decreases with increasing numbers of ipsilateral flankers

Temporal order discrimination improved with increasing temporal separation between the auditory reference and the target tactile landmark across all conditions, but performance progressively decreased as the number of ipsilateral flankers increased (Figure 1C). In the absence of flankers, average performance increased from 59.0*±*2.7% (SE across participants) at 50 ms to 72.0*±*3.7% at 150 ms and 83.2*±*3.9% at 300 ms. Introducing a single ipsilateral flanker reduced performance by a nearly constant factor across temporal separations, with overall performance reaching 0.881*±*0.018 times that of the no-flanker condition (50 ms: 0.875*±*0.051; 150 ms: 0.895*±*0.036; 300 ms: 0.884*±*0.022). The addition of a second ipsilateral flanker produced a further reduction, with overall performance decreasing to 0.857*±*0.017 times that of the no-flanker condition.

To quantify the extent to which ipsilateral flankers affected the temporal precision with which participants discriminated the target tactile landmark, we fit a hierarchical trial-level generalised linear mixed-effects model with a binomial response distribution and logit link. Accuracy was modelled as a function of the absolute temporal separation between the auditory reference and target tactile landmark, with flanker condition included as a modulation of temporal sensitivity and participant-specific random slopes for temporal separation. The model was parameterised without an intercept, constraining performance to chance level at zero temporal separation. Under this formulation, the logit slope for temporal separation quantifies temporal precision, with steeper psychometric slopes corresponding to more precise temporal discrimination.

Including flanker condition significantly improved model fit compared with a null model containing temporal separation alone (ΔAIC = *−*43.8, ΔBIC = *−*23.7 likelihood-ratio test: *χ*^2^(3) = 49.82, *p <* 10*^−^*^10^). The model predicted empirical accuracies with a root mean squared error (RMSE) of 0.058 (5.8 percentage points on the accuracy scale, also see Figure 1C) and showed no evidence of overdispersion (dispersion parameter = 1.00).

Discrimination accuracy increased significantly with temporal separation (*β* = 5.98 *±* 0.75 SE, 95% CI [4.52, 7.44], *t*(5996) = 8.02, *p <* 10*−*15). Relative to the no-flanker condition, a single ipsilateral distractor significantly reduced temporal sensitivity (Δ*β* = *−*2.32 *±* 0.46 SE, 95% CI [*−*3.22*, −*1.42], *t*(5996) = *−*5.08, *p <* 10*^−^*^6^), and two ipsilateral flankers produced a further reduction (Δ*β* = *−*3.03 *±* 0.45 SE, 95% CI [*−*3.92*, −*2.15], *t*(5996) = *−*6.73, *p <* 10*^−^*^11^). Temporal sensitivity showed a progressive decline from *β* = 5.98 *±* 0.75 s^-1^ in the no-flanker condition to 3.66*±*0.72 s^-1^ with one flanker, and 2.95*±*0.72 s^-1^ with two flankers (Figure 1D).

To quantify the extent to which flanker number influenced temporal uncertainty and perceived timing, trial-level choice responses were analysed using a hierarchical generalised linear mixed-effects model with a binomial response distribution and probit link (Figure 2). Candidate models were different in whether flanker condition modulated only the psychometric slope or both the slope and intercept, hence, unlike the accuracy analysis, enabling estimation of temporal uncertainty and the point of subjective synchrony (PSS). Relative to a model containing only temporal offset, allowing flanker-dependent changes in psychometric slope substantially improved model fit (ΔAIC = *−*52.6, ΔBIC = *−*26.7, likelihood-ratio test: *χ*^2^(6) = 64.62, *p* = 5.15 *×* 10*^−^*^12^). Extending this model to include flanker-specific intercepts produced a further improvement in fit (ΔAIC = *−*5.8, ΔBIC = +14.3, likelihood-ratio test: *χ*^2^(3) = 11.79, *p* = 0.008). The full model contained a significant positive common intercept, corresponding to an overall bias towards target-first responses (*β* = 0.12 *±* 0.06 SE, 95% CI [0.01, 0.23], *t*(5992) = 2.07, *p* = 0.039). However, the flanker-specific intercept terms were small and not significantly different from zero for either the one-flanker (*β* = *−*0.07 *±* 0.05 SE, 95% CI [*−*0.17, 0.02], *t*(5992) = *−*1.49, *p* = 0.14) or two-flanker (*β* = 0.01 *±* 0.05 SE, 95% CI [*−*0.08, 0.11], *t*(5992) = 0.26, *p* = 0.79) conditions. Model diagnostics indicated no evidence of overdispersion (dispersion parameter = 1.00). As the flanker-specific intercepts contributed little to the psychometric functions for the no-, one-, and two-flanker conditions, Figure 2A shows predictions from the slope-only model, which produced virtually identical fits to the data (RMSE = 0.10 for both models).

**Figure 2.**
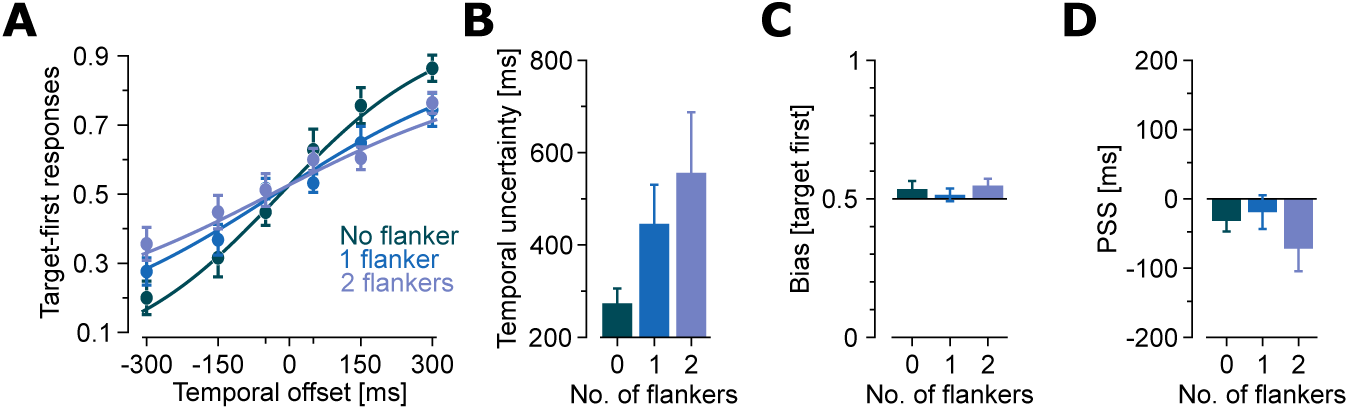
Effect of number of flankers on temporal order judgement, temporal uncertainty, and perceived timing. **A**, Psychometric functions showing the proportion of target-leading responses as a function of the target–auditory reference temporal offset for the no-flanker, one-flanker, and two-flanker conditions. Markers represent the empirical mean proportion of target-leading responses across participants. Error bars denote SE across participants. Solid curves represent the population-level predictions of the hierarchical generalised linear mixed-effects model averaged across subject-specific predictions. **B**, Model-derived estimates of temporal uncertainty for each flanker condition. Error bars represent model-estimated SE. **C**, Empirical response bias, quantified as the net proportion of target-leading responses averaged across all temporal offsets. Error bars denote SE across participants. **D**, Point of subjective synchrony (PSS) estimated from the hierarchical generalised linear mixed-effects model. Error bars represent model-estimated SE.

Consistent with the accuracy analysis (Figure 1C and D), the fitted psychometric slopes decreased progressively with increasing flanker number, corresponding to a monotonic increase in temporal uncertainty (Figure 2B). The estimated temporal uncertainty increased from 273*±*32 ms in the no-flanker condition to 446*±*85 ms with one flanker and 556*±*131 ms with two flankers.

### Number of flankers had little influence on perceived event timing

Because target-leading and target-lagging trials were presented with equal probability, the overall proportion of target-first responses provides a direct behavioural measure of temporal bias. Participants reported the target as occurring first on 53.5 *±* 2.9% of trials in the no-flanker condition, 51.4 *±* 2.3% with one flanker, and 54.7 *±* 2.5% with two flankers (Figure 2C).

Consistently, The model-derived PSS estimates were also consistently negative, indicating that the target vibration had to precede the auditory reference, on average, to be perceived as synchronous. Although the flanker-specific intercept terms were not significantly different from zero, the estimated PSS varied across conditions because the common intercept remained positive while the psychometric slopes differed. Consequently, the estimated PSS was *−*32 *±* 15 ms in the no-flanker condition, *−*19 *±* 24 ms with one flanker, and *−*72 *±* 33 ms with two flankers (Figure 2D).

Together, these results indicate that increasing the number of flankers primarily reduced temporal precision, whereas the changes in perceived event timing were comparatively small and were not explained by flanker-specific shifts of the psychometric function.

### Contralateral flanker decreases temporal precision irrespective of spatial proximity

To quantify the extent to which the reduction in temporal precision depended on spatial proximity, the contralateral flanker condition was compared with the no-flanker and one-flanker conditions (Figure 3). Relative to the no-flanker condition, the contralateral flanker decreased mean discrimination accuracy across temporal separations by 8.3 *±* 2.4% (mean *±* SE across participants, Figure 3A), comparable to the reduction produced by a single ipsilateral flanker (11.9 *±* 1.8%). The hierarchical generalised linear mixed-effects model likewise showed a significant reduction in temporal sensitivity for the contralateral flanker relative to the no-flanker condition (Δ*β* = *−*1.83 *±* 0.46, 95% CI [*−*2.74*, −*0.93], *t*(5996) = *−*3.97, *p* = 7.2 *×* 10*^−^*^5^). Re-referencing the model to the contralateral condition revealed no significant difference in temporal sensitivity between the contralateral and one-flanker conditions (Δ*β* = *−*0.49 *±* 0.43, 95% CI [*−*1.32, 0.35], *t*(5996) = *−*1.14, *p* = 0.26), whereas the two-flanker condition exhibited lower temporal sensitivity than the contralateral condition (Δ*β* = *−*1.20 *±* 0.42, 95% CI [*−*2.02*, −*0.37], *t*(5996) = *−*2.85, *p* = 0.0045).

**Figure 3.**
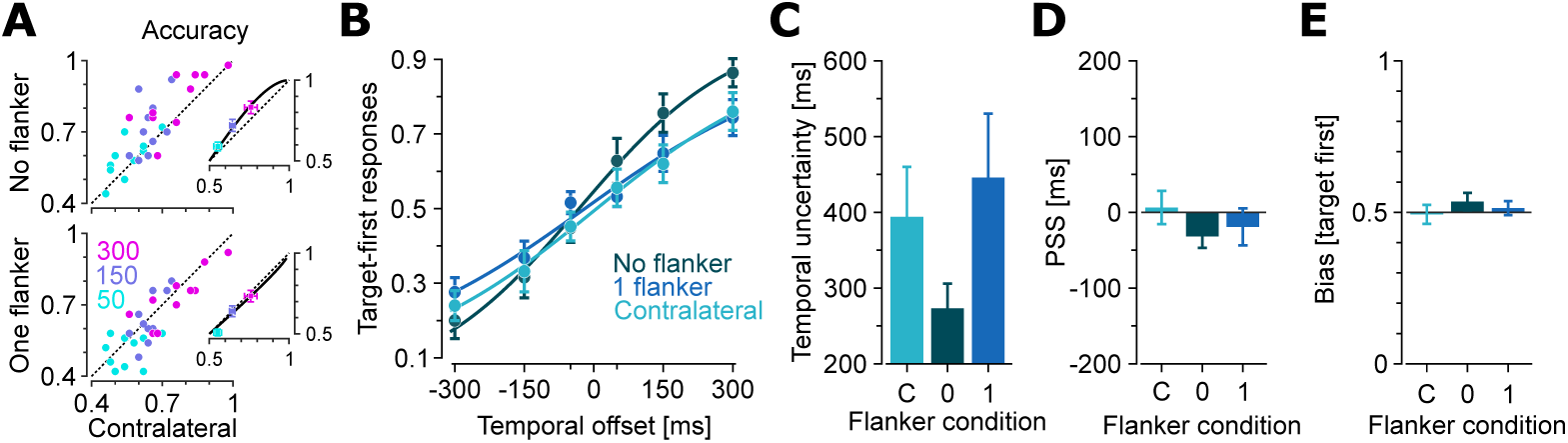
Effect of contralateral flanker on temporal precision, temporal uncertainty, and perceived timing. **A**, Temporal order discrimination accuracy for each participant and temporal separation condition in the no-flanker (top) and one-flanker (bottom) conditions plotted against the corresponding contralateral flanker condition. Insets show cross-participant averages. Error bars denote SE across participants. Solid curves represent population-level predictions from the hierarchical generalised linear mixed-effects model obtained by averaging participant-specific predictions. Dotted lines indicate the line of identity. **B**, Psychometric functions showing the proportion of target-first responses as a function of the target–auditory reference temporal offset for the no-flanker, one-flanker, and contralateral flanker conditions. Markers represent the empirical mean across participants. Error bars denote SE across participants. Solid curves show the population-level predictions of the hierarchical generalised linear mixed-effects model. **C**, Model-derived temporal uncertainty for the contralateral (C), no-flanker (0), and one-flanker (1) conditions. Error bars represent model-estimated SE. **D**, Point of subjective synchrony (PSS) estimated from the hierarchical generalised linear mixed-effects model. Error bars represent model-estimated SE. **D**, Behavioural bias quantified as the overall proportion of target-first responses across temporal offsets. Error bars denote SE across participants.

The psychometric functions exhibited a similar reduction in slope for the contralateral and one-flanker conditions (Figure 3B). Temporal uncertainty increased from 273 *±* 32 ms in the no-flanker condition to 394 *±* 66 ms in the contralateral condition, compared with 446 *±* 85 ms for one ipsilateral flanker (Figure 3C).

In contrast to temporal precision, the contralateral flanker produced little effect on perceived event timing. The estimated PSS was close to zero for the contralateral condition (6 *±* 22 ms), compared with *−*32 *±* 15 ms and *−*19 *±* 24 ms for the no-flanker and one-flanker conditions, respectively (Figure 3D). The direct behavioural measure of bias showed the same pattern. Participants reported the target as occurring first on 49.3 *±* 3.1% of trials in the contralateral flanker condition, compared with 53.5 *±* 2.9% and 51.4 *±* 2.3% in the no-flanker and one-flanker conditions, respectively (Figure 3E).

Together, these findings indicate that a flanker presented on the opposite hand degrades temporal precision almost as strongly as a single ipsilateral flanker while producing little change in perceived event timing. The comparable effects of contralateral and ipsilateral flankers indicate that a substantial component of the temporal interference is independent of local spatial proximity, consistent with a central mechanism rather than one arising solely from interactions between neighbouring tactile stimuli.

### Relative flanker timing biases perceived temporal order without affecting temporal precision

The analyses so far quantify the spatial contextual effect revealing that increasing the number of flankers reduced discrimination accuracy and temporal sensitivity irrespective of flanker location, without producing a systematic change in perceptual bias and the point of subjective simultaneity. How does the temporal context influence temporal precision and perceived timing? In this paradigm, each flanker reached its envelope peak either 60° before or 60° after the target peak (*±*333 ms; Figure 1B), allowing the effects of leading and lagging tactile context on temporal sensitivity and perceived timing to be compared directly.

Adding flanker timing to the accuracy GLMM showed no significant change in temporal sensitivity after accounting for flanker condition (*β* = 0.12 *±* 0.19, 95% CI [*−*0.49, 0.25], *t*(5995) = *−*0.64, *p* = 0.523). The model including the effect of flanker timing on the psychometric slope also did not fit the data better than the model containing flanker condition alone (ΔAIC = +1.6, ΔBIC = +8.2, likelihood-ratio test: *χ*^2^(1) = 0.41, *p* = 0.523). Little change in sensitivity was also evident in the empirical data, with accuracy for leading and lagging flankers remaining close to the identity line across temporal separations and across the one-flanker, two-flanker, and contralateral conditions, as shown in Figure 4A.

**Figure 4.**
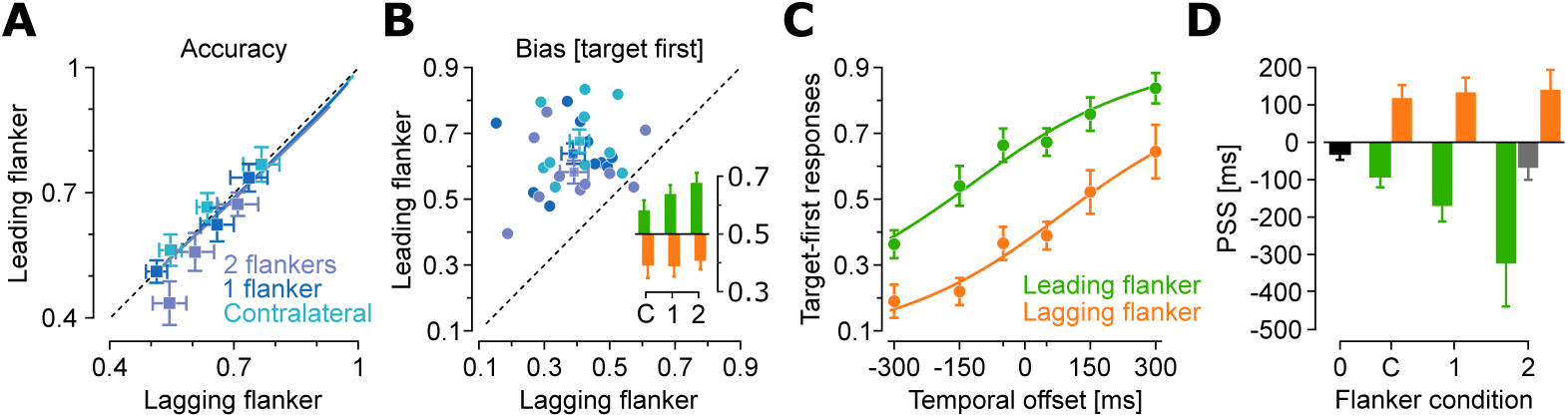
Relative flanker timing shifts perceived temporal order without altering temporal precision. **A**, Temporal order discrimination accuracy for each temporal separation condition for leading versus lagging flanker conditions. Data are shown separately for the one-flanker, two-flanker, and contralateral flanker conditions. Markers represent the mean proportion of correct responses across participants and error bars denote SE. Solid curves show the population-level predictions of the hierarchical generalised linear mixed-effects model averaged across participants. The dashed line indicate the line of identity. **B**, Behavioural bias, expressed as the proportion of target-first responses, for leading and lagging flanker conditions. Square markers represent participant means and circles denote individual participants for the one-flanker, two-flanker, and contralateral conditions. Colour conversion is as in**A**. Inset: mean behavioural bias for leading (green) and lagging (amber) flanker conditions. Error bars denote SE across participants. **C**, Psychometric functions for the one-flanker condition showing the proportion of target-first responses as a function of the target–auditory reference temporal offset for leading and lagging flankers. Markers represent the empirical mean across participants. Error bars denote SE across participants. Solid curves show the population-level predictions of the hierarchical generalised linear mixed-effects model averaged across participants. **D**, Point of subjective synchrony (PSS) estimated from the psychometric GLMM. The no-flanker condition is shown as a single reference bar (black). For the two-flanker condition, the grey bar represents the balanced condition in which one flanker led and the other lagged the target. Error bars denote model-estimated SE.

In contrast, perceptual bias showed a different pattern. Participants reported that the target occurred first more frequently when the flankers led the target than when they lagged behind it (Figure 4B). The effect was markably robust at individual level and across all flanker configurations, including the contralateral condition where the mean percentage of target-first responses increased from 39.1*±*4.4% for lagging flankers to 58.2*±*3.5% for leading flankers. Similar pattern was observed for one ipsilateral flanker, increasing from 38.8*±*3.6% to 63.8*±*3.1%, and for two flankers, increasing from 40.7*±*3.0% when both flankers lagged the target to 67.6*±*3.5% when both led it. The direction of these shifts indicates that the perceived timing of the target peak was attracted towards the timing of the flankers.

### Leading and lagging flankers produce opposite temporal biases

The observed pattern of flanker timing was captured by psychometric analysis and trial-level hierarchical GLMM of target-first responses with a binomial distribution and logit link Figure 4C. Consistent with the accuracy analysis, allowing the psychometric slope to vary with flanker timing did not provide evidence for an additional change in temporal sensitivity. The effect of flanker timing was instead a systematic shift of the psychometric intercept, corresponding to horizontal shifts of the psychometric functions along the temporal-offset axis (Figure 4C). Relative to the no-flanker condition as a reference (intercept = 0.19 *±* 0.10, 95% CI [0.00, 0.38], *t*(5984) = 2.01, *p* = 0.044), leading flankers exhibited significant positive shifts (contralateral: 0.21 *±* 0.10, 95% CI [0.02, 0.41], *t*(5984) = 2.16, *p* = 0.031; one-flanker: 0.48 *±* 0.10, 95% CI [0.28, 0.68], *t*(5984) = 4.68, *p <* 10*^−^*^5^, two-flankers: 0.66 *±* 0.13, 95% CI [0.41, 0.91], *t*(5984) = 5.12, *p <* 10*^−^*^6^), corresponding to higher proportion of target-first responses. In contrast, lagging flankers exhibited significant negative shifts (contralateral: *−*0.71 *±* 0.10, 95% CI [*−*0.91*, −*0.50], *t*(5984) = *−*6.81, *p <* 10*^−^*^10^; one-flanker: *−*0.73 *±* 0.10, 95% CI [*−*0.93*, −*0.53], *t*(5984) = *−*7.17, *p <* 10*^−^*^12^, two-flankers: *−*0.67 *±* 0.13, 95% CI [*−*0.92*, −*0.42], *t*(5984) = *−*5.25, *p <* 10*^−^*^6^), corresponding to lower proportion of target-first responses.

As the PSS is defined by the psychometric intercept relative to its slope, these intercept shifts translated directly into systematic changes in the estimated PSS (Figure 4D). Consistent with the perceptual bias measured directly from the responses (Figure 4B), when the flankers led the target, the estimated PSS was negative for the contralateral, one-flanker, and two-flanker conditions (*−*92 *±* 28, *−*170 *±* 41, and *−*324 *±* 115 ms, respectively), indicating that the target was perceived earlier relative to the auditory reference. When the flankers lagged the target, the estimated PSS shifted in the opposite direction to 118 *±* 35, 133 *±* 40, and 140 *±* 54 ms, respectively), indicating that the target had to occur later relative to the auditory reference to be perceived as simultaneous.

Together, the psychometric analysis, PSS estimates, and direct behavioural bias provide consistent evidence that leading flankers shift the perceived timing of the target towards earlier times, whereas lagging flankers shift it towards later times, while temporal sensitivity remains unchanged.

### Opposing flanker timing generates motion-dependent temporal bias

For the two-flanker condition, the flankers were assigned independently to lead or lag the target. Consequently, the mixed condition, in which one flanker preceded and the other followed the target, produced an intermediate behavioural bias (55.2 *±* 3.2%) close to the no-flanker condition (53.5 *±* 2.9%), together with an intermediate PSS (*−*68 *±* 33 ms versus *−*32 *±* 16 ms for the no-flanker condition, Figure 4D). At first glance, this may suggests that the opposing temporal influences of the two flankers largely cancel.

However, opposite flanker timings also define opposite directions of apparent tactile motion across the three fingers [45], as shown in Figure 5A. We therefore asked whether the mixed condition contains two distinct perceptual states depending on the direction of apparent motion rather than simply averaging to zero. The mixed condition was further separated according to the direction of apparent tactile motion. Motion from the index towards the ring finger (radial-to-ulnar direction) produced a larger proportion of target-first responses (59.4*±*2.7%) than motion in the opposite direction (50.7*±*4.5%; Wilcoxon signed-rank test, *W* = 47, *p* = 0.049, Figure 5B and C). These results suggest that even when the overall flanker lead and lag influences are balanced, the direction of apparent tactile motion continues to bias perceived temporal order.

**Figure 5.**
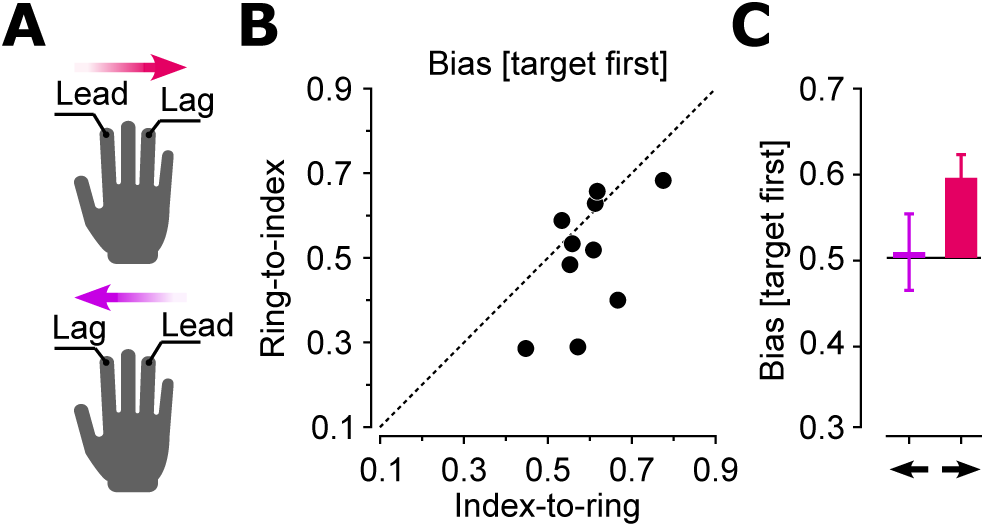
Opposing flanker timing generates motion-dependent temporal bias. **A**, Schematic of the mixed two-flanker condition. One flanker led and the other lagged the target by 60° of envelope phase, producing two opposing apparent motion sequences across the three fingers: index-to-ring (radial-to-ulnar) and ring-to-index (ulnar-to-radial). **B**, Perceptual bias, expressed as the proportion of target-first responses for the index-to-ring motion condition (abscissa) versus the ring-to-index motion condition (ordinate). Each marker represents one participant. The dashed line indicates the line of identity. **C**, Mean proportion of target-first responses for the two motion directions. Error bars denote SE across participants.

## Discussion

Increasing the number of flankers progressively reduced temporal precision, increasing perceptual uncertainty. A contralateral flanker produced a reduction of comparable magnitude to that produced by a single adjacent ipsilateral flanker, indicating that this interference was not restricted to neighbouring fingers or local spatial proximity.

Relative flanker timing produced a different effect: leading flankers shifted the perceived timing of the target earlier, whereas lagging flankers shifted it later, without changing temporal precision. When one flanker led and the other lagged, the perceptual bias differed between the two resulting across-finger sequences, with radial-to-ulnar motion producing more target-first reports than ulnar-to-radial motion. Together, these findings distinguish contextual effects on the uncertainty of a tactile temporal estimate from contextual shifts in the estimated time of tactile representations.

### Distributed tactile context reduces temporal precision

The progressive reduction in temporal precision with increasing flanker number extends earlier work on tactile masking and temporal interference [35, 38, 64–66]. Previous studies showed that a temporally adjacent nontarget can impair detection of target stimuli, or geometric identification of tactile patterns through both degradation of the target representation and competition between target- and nontarget-related responses [40, 65, 66]. Similar interference has been reported when concurrent stimulation was delivered to different fingers, indicating that tactile information cannot always be restricted to the task-relevant location [40, 67]. The present results extend these findings from feature identification and detection to the precision with which the time of an internal tactile landmark is represented. The monotonic increase in temporal uncertainty with one and two flankers suggests that additional tactile inputs reduce the reliability of the target’s temporal estimate rather than producing only a fixed response bias. Multi-digit responses and nonlinear interactions across fingers have been identified in human somatosensory cortex [39, 68, 69], providing plausible neural substrates for the progressive loss of temporal precision as additional ipsilateral flankers were introduced.

The effect of the contralateral flanker further indicates that this observed reduced precision is not restricted to local interactions between adjacent fingers. Long-range tactile masking has previously been observed across homologous regions of the two sides of the body [70], and a recent study using a transient target embedded within an ongoing vibration found robust interference in target detectability from stimulation of homologous and non-homologous fingers on the opposite hand [71]. Interference showed small differences in finger identity and hand separation, and target detection reflected combined information from the attended and unattended sides [71]. The present finding that an opposite-hand flanker reduced temporal precision by an amount similar to an adjacent flanker is consistent with such distributed interactions. This result does not establish equivalent effects across ipsilateral and contralateral conditions, because these conditions were not tested for statistical equivalence. It instead argues against an account based solely on overlap between neighbouring peripheral or cortical finger representations. Tactile inputs from the two hands interact within somatosensory cortex, including the primary and secondary somatosensory cortices (S1 and S2) in both humans and rodents [72–76], providing a plausible substrate for cross-hand interference.

### Temporal context shifts perceived timing

Unlike the reduction in temporal precision produced by additional flankers, changes in flanker timing displaced the temporal estimate without reducing its precision. Leading flankers shifted the target towards earlier perceived times, whereas lagging flankers shifted it later. This pattern differs from a general masking account in which temporally adjacent stimulation weakens the target representation. Instead, the target estimate was shifted towards the temporal context while the temporal sensitivity remained stable. Attractive temporal biases have been reported when the timing or trajectory of task-irrelevant sensory information is consistent with a broader event sequence, although most tactile TOJ studies have examined the order of discrete stimulus onsets rather than the timing of an internal feature within an extended signal [55, 58, 77, 78].

The direction of the shift is compatible with temporal integration of target and flanker information. Earlier tactile masking studies showed that nearby patterns can be combined into composite percepts and that temporal integration and response competition can occur under the same stimulus conditions [40, 66]. A recent study similarly found that concurrent tactile signals can be pooled across the hands even when one site is explicitly irrelevant [71]. The observed temporal shifts do not necessarily imply pooling of the spatial identities of the target and flankers. Instead, they suggest that temporal context displaced the temporal estimate of the target landmark towards the temporal structure of the surrounding tactile signals. This interpretation is consistent with the absence of a measurable change in temporal precision, indicating that the temporal estimate was systematically shifted without a corresponding increase in temporal uncertainty.

### Dissociation between temporal precision and perceived timing

The dissociation between flanker number and flanker timing shows that contextual stimulation does not operate as a single source of undifferentiated temporal noise. Adding flankers reduced the reliability of the temporal estimate, whereas changing their relative timing displaced its central timing value. These effects correspond to separable properties of the estimated temporal representation: its uncertainty and its perceived location in time.

Changes in temporal-order judgements can arise from multiple sources, including increased sensory uncertainty, shifts in perceived event timing, spatial remapping, changes in decision criteria, or interactions among these processes [55, 57, 79]. The present results distinguish two of these possibilities. Increasing surrounding tactile input reduced temporal precision, whereas changing its temporal relationship to the target systematically shifted perceived timing without reducing precision. Neural studies likewise distinguish processes contributing to temporal sensitivity from those involved in temporal ordering, spatial remapping, and simultaneity judgements [57, 79, 80]. Any account of tactile landmark timing must therefore explain how increasing surrounding tactile input broadens the temporal estimate, whereas changing the timing of that input systematically shifts the estimate without increasing its uncertainty.

### Motion-dependent bias

The mixed lead–lag condition also revealed a sequence-dependent bias. Radial-to-ulnar motion produced more target-first reports than the reverse sequence, even though the number of leading and lagging flankers were matched. Previous studies provided a close precedent that task-irrelevant within-finger motion of tactile patterns biased order judgements [58, 77]. When the first pattern moved towards the location of the second, participants were more likely to judge the first stimulus as leading, while motion in the opposite direction produced the reverse tendency. A cross-modal study showed that the spatial relationship between visual and tactile motion trajectories altered TOJs and disappeared when the displays were repositioned so that the trajectories no longer converged [78]. These studies show that motion structure can alter temporal order even when motion is irrelevant to the task.

The present result extends this principle to motion generated by the relative phase of signals across fingers rather than by lateral movement within each stimulus. Relative envelope phase is sufficient to produce apparent tactile motion across stationary fingerpads [45, 46, 50], and the sequence-dependent bias is therefore compatible with motion influencing the temporal estimate of the central landmark. However, apparent-motion direction was confounded with the identity of the leading flanker (index versus ring fingerpads). A recent multidigit study showed that within-hand integration associates greater perceptual weight to the finger leading a tactile trajectory [42]. The difference between radial-to-ulnar and ulnar-to-radial sequences could therefore reflect an extracted motion signal, unequal weighting of index and ring finger inputs, or both. The present data support a sequence-dependent bias but do not separate these explanations. Future experiments that independently manipulate motion direction and finger identity will be required to dissociate these possibilities.

### Computational mechanisms

Two broad accounts can explain this pattern. Under a response-competition account, target and flanker signals have separate representations but contribute competing evidence to the final temporal decision [40, 65, 81]. Increasing the number of flankers would increase the probability that the judgement is influenced by temporal evidence originating from an irrelevant tactile signal, thereby reducing temporal precision. Leading and lagging flankers would bias the accumulated evidence towards earlier or later target estimates. This account can therefore explain both the reduction in temporal precision and the systematic shifts in perceived timing.

Under a pooling account, target and flanker signals are combined before the temporal decision into a weighted temporal representation. Additional flankers would broaden or destabilise this representation, reducing temporal precision, whereas leading or lagging flankers would shift it towards earlier or later times. Evidence for obligatory or near-obligatory integration across tactile sites comes from multi-digit motion averaging, nonlinear Pooling naturally predicts the present dissociation because increasing the number of contributing signals broadens the temporal estimate, whereas changing their relative timing displaces the estimate without necessarily increasing its uncertainty. Behavioural evidence for obligatory or near-obligatory integration across tactile sites comes from multi-digit motion integration [41, 42]. Physiological studies likewise reveal nonlinear interactions across multiple fingers in human somatosensory cortex [68, 69], while behavioural interactions also extend across the two hands during tactile detection [71]. Human somatosensory cortex additionally exhibits normalisation and feature-dependent interactions during concurrent tactile stimulation [39, 82]. These findings make pooling a plausible account, although the present behavioural data do not identify the processing stage at which signals are combined.

The two accounts are not mutually exclusive. Earlier work concluded that temporal integration and response competition coexist during tactile pattern perception [66, 81]. A pooled sensory representation could therefore be followed by competition among target- and flanker-related evidence during the decision stage. Distinguishing these possibilities will require manipulations that independently vary sensory pattern, task relevance, and response competition. For example, changing flanker waveform while preserving landmark timing would quantify the contribution of sensory pattern, whereas independently manipulating motion direction and leading-finger identity would distinguish motion-based pooling from unequal weighting across fingers.

### Link to flash-lag effect and motion-induced mislocalisation

The present findings shares conceptual similarities to the flash-lag family of perceptual phenomena. Rather than comparing the position of a moving object with a flash [83–86], participants compared the timing of a salient landmark embedded within a continuously evolving tactile envelope against a brief auditory reference. Similar extensions of the flash-lag paradigm have shown that the effect is not restricted to spatial position, but also occurs for continuously changing stimulus attributes including colour, luminance, orientation and other feature dimensions [87–89]. The present task therefore extends this continuous-feature versus transient-reference comparison to vibrotactile magnitude and landmark timing. The small baseline PSS should not be interpreted directly as evidence for or against a tactile flash-lag effect because it includes modality-specific delays between tactile and auditory processing. Establishing a genuine tactile feature flash-lag would require comparison with a transient tactile reference. Nevertheless, the flash-lag literature provides an informative comparison because several paradigms show that the direction and magnitude of the effect depend strongly on changes in the temporal evolution of the continuous stimulus. Motion reversal and abrupt termination can markedly reduce or even reverse the flash-lag effect [84–86, 89, 90], indicating that the perceived state of a continuously evolving stimulus depends on how its temporal evolution is interpreted rather than on a simple forward extrapolation of its trajectory.

A second connection arises as the target and flankers generated tactile motion across the fingers due to their phase-offset [45]. With one flanker, the direction of apparent motion depended on whether the flanker preceded or followed the target, producing trajectories from the index to the middle finger or from the middle to the index finger. Importantly, the auditory probe occurred while the target envelope continued to evolve for approximately 700–1,300 ms after the probe, such that the perceived illusory tactile source described in our previous work [45] remained dynamic after the temporal comparison was made. This resembles flash-lag paradigms in which a continuously moving object is compared with a transient flash, except that here the moving percept is generated by the evolving tactile envelopes rather than by physical displacement across space. The direction-dependent shifts observed here therefore raise an interesting possibility. Rather than assigning time to the physical peak of the target envelope, marking the position of the illusory tactile source under at the fingertip, the nervous system may assign time to the perceived position of the evolving tactile object generated by the target and flankers. In this interpretation, leading flankers would advance the apparent position of the moving tactile percept in time, whereas lagging flankers would delay it, analogous to motion-induced shifts in perceived position reported in the visual flash-lag literature [86, 89, 91, 92]. The mixed two-flanker condition is particularly informative because the target formed the central element of a continuous apparent-motion trajectory, yet opposite motion directions still produced different temporal estimates. Whether this reflects temporal localisation of the moving tactile percept itself, or unequal weighting of the leading finger, remains unresolved.

### Concluding remarks

The present study used an auditory reference to estimate the timing of the tactile landmark. Although the measured point of subjective simultaneity incorporates modality-specific processing delays [93–95], these delays were constant across conditions and therefore cannot account for the observed contextual shifts. Auditory timing provides a temporally precise external reference, making it well suited for isolating tactile contextual effects. Nevertheless, future studies using tactile reference stimuli will establish the extent to which the present findings generalise within the tactile modality.

The present findings show that tactile temporal perception is influenced not only by the presence of surrounding tactile signals but also by their spatiotemporal organisation. Distributed tactile context reduced the precision of temporal estimates, whereas temporal context systematically displaced perceived event timing. These results extend contextual interactions in touch from feature identification and onset order to the temporal representation of tactile landmarks embedded within continuous signals, providing a foundation for investigating how distributed tactile inputs are integrated during natural haptic perception.

## Notes

### Competing Interest Statement

The authors have declared no competing interest.

### Funding

- Australian Research Council: DE200101468

